# Naturally occurring ACE2 stalk variants are differentially released from the cell

**DOI:** 10.64898/2026.04.27.721131

**Authors:** Florian Wiersch, Christine Lux, Julia Vanderliek-Kox, Katharina Schun, Andreas Ludwig, Stefan Düsterhöft

## Abstract

Angiotensin-converting enzyme 2 (ACE2) is a key regulator of the renin-angiotensin-aldosterone system (RAAS). It also acts as a receptor for SARS-CoV-2 and stabilises the B0AT1 amino acid transporter at the cell surface. Therefore, surface expression of ACE2 is crucial for these physiological processes. ACE2 is released as a soluble, catalytically active form, partly through ectodomain shedding. This process mainly involves the sheddases ADAM10 and ADAM17, but the exact regulatory mechanisms remain unclear.

We assessed 11 naturally occurring single-point mutations in the ACE2 stalk region. Most variants showed significantly reduced release compared to wild-type (WT) ACE2; however, the single point mutations P734L and G726R significantly increased their release. ACE2_P734L also exhibits higher surface expression, directly increasing the surface levels of B0AT1. Despite B0AT1 and ACE2 forming a tight tetrameric complex, this did not affect ACE2 shedding. This suggests that complex formation does not restrict sheddase access.

Overall, these data identify the ACE2 stalk region as a major determinant of shedding efficiency. Naturally occurring variants in this region can substantially affect the release of soluble ACE2, potentially contributing to interindividual differences that are relevant for pathophysiological processes.

## Introduction

Angiotensin-converting enzyme 2 (ACE2) is involved in numerous (patho)physiological processes due to its crucial role within the renin-angiotensin-aldosterone system (RAAS). RAAS is activated upon decreased renal perfusion resulting in release of renin, which consecutively cleaves angiotensinogen to angiotensin I in the bloodstream. Angiotensin converting enzyme 1 (ACE1), a homologue of ACE2, converts angiotensin I to angiotensin II (Ang II). Ang II mediates retention of water and salt, vasoconstriction as well as inflammatory, fibrotic and prothrombotic stimuli to local tissue via its receptor AT1 (Donoghue et al., 2000, Rice et al., 2004). The ability of ACE2 to process Ang II to Angiotensin 1-7 (Ang-(1-7)), a peptide exerting tissue protective effects via its receptor Mas, makes it a key regulatory point within the RAAS. This is emphasized by the fact that ACE2 was found to be protective in several cardiovascular diseases such as heart failure, hypertension and renal fibrosis in ACE2 deficiency mouse models (Jia et al., 2020). Another major function of ACE2 is its role in cellular uptake of several coronaviruses such as SARS-CoV-2, which binds to ACE2 via its spike protein prior to endocytosis (Li et al., 2003, Cui et al., 2019, Hoffmann et al., 2020). Because of its viral receptor function and also because of its protective activity in RAAS-mediated tissue damage, ACE2 is of great importance for SARS-CoV-2 infectivity and COVID-19 disease (Elshafei et al., 2021, Rysz et al., 2021). Several risk factors such as old age, male sex and cardiovascular comorbidities as well as several gene polymorphisms within the ACE2 gene are known to be associated with a severe clinical course of COVID-19 (Zhang et al., 2023).

ACE2 evolved through fusion of TMEM27 (also known as collectrin) and ACE1, explaining its extracellular structure comprising a metalloprotease domain (MD) and a collectrin-like domain (CLD) (Zhang et al., 2001). Because of their homology, TMEM27, ACE1 and ACE2 can be grouped together as part of the ACE2 family (Webers et al., 2024). The extracellular domains of ACE2 (MD and CLD) are linked to the transmembrane helix by the stalk region consisting of a flexible and unstructured chain of 21 amino acids (Niehues et al., 2022).

ACE2 is predominantly expressed as a membrane-bound peptidase (mACE2) in the intestine, kidney, testis, gallbladder, heart, and to a lesser extent also in the lung (Hikmet et al., 2020). Membrane anchoring restricts the activity of ACE2, and thus the local degradation of Ang II and the generation of Ang-(1–7), to tissues and cells that express it. This protective function can be reduced by the proteolytic cleavage of the ACE2 ectodomain (shedding), which removes ACE2 from the cell surface (Lambert et al., 2005, Niehues et al., 2022). The resulting soluble ectodomain (sACE2) retains its catalytic activity, enabling ACE2 to exert its effects in the extracellular environment, including in tissues where membrane-bound ACE2 expression is low. ACE2 can also be removed from the cell in a soluble form by extracellular vesicles (Niehues et al., 2022). Both mechanisms can lead to reduction of ACE2 surface expression which may critically impact ACE2-mediated SARS-CoV-1/2 infectivity and cardioprotective functions (Li et al., 2003, Cao et al., 2020).

Shedding of ACE2 is predominantly mediated by ADAM17 and ADAM10, both belonging to the ADAM (a disintegrin and metalloproteinase) family (Lambert et al., 2005, Niehues et al., 2022, Webers et al., 2024). Noteworthy, constitutive ACE2 shedding primarily involves ADAM10 while ADAM17 seems to be responsible for induced shedding (Niehues et al., 2022). The release of the other members of ACE2 family, TMEM27 and ACE1, is partially mediated by different sheddases. While ACE1 is predominantly shed by ADAM10, TMEM27 is mainly shed by BACE2 (Webers et al., 2024). Notably, ADAM17 and ADAM10 do not have universal recognition sequences, yet there are some patterns occurring frequently in cleavage sites of their substrates (Tucher et al., 2014). Cleavage sites of most substrates are located in the flexible juxtamembranous stalk region, suggesting that ACE2 stalk region harbours the cleavage site(s). We could previously show that an ACE2 construct consisting only of the collectrin-like parts of ACE2 exhibited similar shedding qualities to full length ACE2, indicating that the metalloprotease domain is dispensable for ACE2 shedding (Niehues et al., 2022).

Besides its role in the RAAS, ACE2 is also responsible for stabilisation of amino acid transporters B0AT1 (SLC6A19) and XTRP3 (SLC6A20) at the cell surface in the intestine. In the kidney meanwhile, this stabilization is provided by collectrin which is not expressed in the intestine (Danilczyk et al., 2006, Camargo et al., 2020). At the cell surface, heterodimers of B0AT1 and ACE2 group together, resulting in a tetramer of two heterodimers (Yan et al., 2020). As this interaction is mediated by the collectrin-like domain of ACE2, it has been suggested that B0AT1 binding physically restricts the access of sheddases to the ACE2 cleavage sites (Andring et al., 2020, Yan et al., 2020, Stevens et al., 2021).

The aim of this study was to characterise the structural determinants of ACE2 shedding. We could show that ACE2 stalk region is one of the key determinants of shedding given that replacement of the stalk region as well as introduction of natural occurring point mutations lead to significant changes in shedding rate. Moreover, we could demonstrate that the ACE2 variant P734L leads to an increased surface expression of B0AT1. However, our data indicate that the presence of B0AT1 does not interfere with the shedding process.

## Material and methods

### 1. Cloning

ACE2 mutants and ACE2 chimeras ACE2_TMEM27, ACE2_ACE1 and ACE2_CD4S were cloned using overlap extension PCR. For detection and activity assays, a 4xmyc tag and/or a human placenta alkaline phosphatase (AP) tag was N-terminally fused to the ACE2 constructs as described before (Niehues et al., 2022, Webers et al., 2024). Analogously, amino acid transporter B0AT1 was C-terminally HA-tagged for Western blot analysis.

### 2. Cell culture, transfection, transduction and cell lysis

HEK293 cells and HEK293 cells deficient for both ADAM17 and ADAM10 (HEK dKO) (Niehues et al., 2022) were cultured in humidified incubators at 37 °C and 5% CO_2_ in DMEM +/+ medium, consisting of DMEM (PAN-Biotech GmbH, Aidenbach, DE) supplemented with 5% FCS (PAN-Biotech GmbH, Aidenbach, DE) and 1% penicillin/streptomycin (Sigma-Aldrich, St. Louis, US). Upon confluency of 70 – 80%, cells were split. HEK293 cells stably overexpressing GFP, B0AT1, ACE2 or ACE2 with the P734L mutation (P734L) were generated through retroviral transduction of HEK293 cells with pMOWS vector system using virus producing Phoenix Ampho cell line (HEK293 cell derivate; ATCC CRL‐3213, ATCC, Manassas, Virginia, USA). Successfully transduced cells carrying zeocin resistance (pMOWS vector system) were selected via antibiotic treatment with zeocin (Invivogen, San Diego, US). For transient overexpression, Lipofectamine 3000 Transfection-Kit (Invitrogen, Thermo Fisher Scientific, Waltham, US) was used according to manufacturer’s manual. For protein biochemistry experiments, transduced or transfected cells were lysed using lysis buffer (50 mM Tris-Base, 150 mM NaCl, 2 mM EDTA, 1% Triton X-100, pH 7.5) supplemented with 10 mM 1,10-phenanthroline (Sigma-Aldrich, St. Louis, US) and 1 tablet cOmplete protease inhibitor (Hoffmann-La Roche AG, Basel, CH) at 4 °C and subsequently centrifuged at 16000xg for 20 minutes. For western blot analysis, 40 µl of obtained supernatant was mixed with 10 µl 5x Laemmli buffer and heated for 20 minutes at 60 °C.

### 3. Supernatant-immunoprecipitation (supernatant-IP) assay

3×10^6^ HEK-293 cells stably overexpressing GFP as control, myc-tagged wt ACE2 or myc-tagged ACE2_P734L were seeded in 20 cm cell culture dishes in DMEM +/+. After 24 hours, medium was changed to OptiMEM containing either 10 µM GI, 40 µM Marimastat or DMSO. After 48 hours of incubation, cells were lysed as described above and supernatants were centrifuged twice (5 minutes at 1000xg followed by 20 minutes at 2880xg) to remove remaining cell (debris). 30 µl magnetic beads diluted in 500 µl lysis buffer were added to the supernatants and samples were incubated rolling for 2 hours at 4 °C. Samples were placed in a magnetic rack and supernatants were discarded. Magnetic beads were transferred to 1.5 ml reaction tubes using 700 µl lysis buffer. A total of 5 washing steps with 700 µl lysis buffer at 4 °C was performed. After removing lysis buffer from the last washing step, 50 µl 2x Laemmli buffer was added. Samples were then heated to 60 °C for 20 minutes for Western blotting analysis.

### 4. Western blotting

After separation via SDS-Page, proteins were transferred onto a methanol activated PVDF-membrane with pore size 0.45 µm (Sigma-Aldrich, St. Louis, US) using Western blotting. Dried and methanol reactivated membranes were blocked for 20 minutes with 5% (w/v) nonfat dry milk in TBS (2 mM Tris HCl, 15 mM NaCl, 1 mM EDTA, pH 7.4) with 1 % Tween-20 (TBST). After three washing steps with TBST, membranes were incubated for 24 hours at 4 °C with primary antibody. Unbound primary antibody was cleared by washing three times with TBST. Then, membranes were incubated with secondary antibody for one hour at room temperature. After one washing step with TBST followed by two washing steps with TBS Odyssey Fluorescence Imager (LI-COR Biosciences GmbH, Lincoln, US) was used for detection of fluorescence-labelled second antibodies. HRP-conjugated antibodies were incubated as described above for secondary antibodies and were visualised with ChemiDoc MP Imaging System (Bio-Rad, Hercules, US) after adding ECL Prime Peroxide Solution (Thermo Fisher Scientific, Waltham, US).

All used antibodies were diluted in TBST + 1% BSA. The following primary antibodies with corresponding dilutions were used: α-myc (1:5000 or 1:2500, ab32, Abcam, Cambridge, UK), α-GAPDH (1:2000 or 1:3333, MA5-15738, Thermo Fisher Scientific, Waltham, US), α-HA (1:1000, 901502, BioLegend, San Diego, US). Goat α-mouse DyLight 680 (1:100,000, 35519, Thermo Fisher Scientific, Waltham, US) was used as secondary antibody. Additionally, HRP-conjugated antibodies α-HA-HRP (1:3333, HAM0601, R&D Systems, Minneapolis, US) and α-myc-HRP (1:2000, ab19312, Abcam, Cambridge, UK) were used. Before use, secondary antibodies were filtered through filters with pore size 0.22 µm. Visualized bands were quantified using Image Studio Lite Version 5.2 (LI-COR Biosciences, Lincoln, US).

### 5. Alkaline phosphatase activity assay

Alkaline phosphatase assay (AP-assay) was used to indirectly analyse release of expressed ACE2 constructs. Cells were seeded in 6-well plates at confluence of 80% and incubated in Opti-MEM (Thermo Fisher Scientific, Waltham, US). 10 µM GI254023X (GlaxoSmithKline, Stevenage, UK), 10 µM GW280264X (Tocris, UK), 10 µM Tapi-1 (Selleck Chemicals GmbH, Planegg, DE) or 10 µM Marimastat (Sigma-Aldrich, St. Louis, US) were used as inhibitors. 0.1 µM PMA was used as a stimulator. For consistency, equal concentration of solvent DMSO was used in all conditions. After 24 hours of incubation, supernatants were harvested and cells were lysed using lysis buffer (pH 7.5, 50 mM Tris-Base, 150 mM NaCl, 2mM EDTA, 1% Triton X-100, 10 mM 1,10-phenanthroline (Sigma-Aldrich, St. Louis, US) and 1 tablet cOmplete protease inhibitor (Hoffmann-La Roche AG, Basel, CH). Measurement and calculation of relative release was done as described before (Niehues et al., 2022, Webers et al., 2024)

### 6. Flow cytometric analysis

HEK293 cells stably expressing wt ACE2 or P734L were transiently transfected with GFP as control or B0AT1 as described above. After detachment using accutase (PAN-Biotech GmbH, Aidenbach, Germany), 2×10^5^ cells were placed in flow cytometry tubes on ice while remaining cells were lysed as described above and served as input control for Western blot analysis. All following steps were conducted on ice or in 4 °C. Flow cytometry tubes were centrifuged (5 minutes at 300x g) and supernatant was discarded. Afterwards cells were washed with 200 µl flow cytometry buffer (FC buffer) consisting of 500 ml sterile PBS and BSA (0,2% w/v and filtered through 0.2 µm pore filters) and centrifugated (5 minutes at 300x g). Supernatant was discarded. Cell pellets were suspended in 45 µl primary antibody and incubated for 1 hour in the dark. After two further washing steps with FC buffer as described above, cell pellet was resolved in 45 µl secondary antibody and left for 45 minutes for incubation. Unbound secondary antibody was removed by two washing steps as described above (first with FC buffer, second with PBS). Resulting cell pellet was resuspended in 400 µl PBS and subjected to flow cytometry analysis using LSRFortessa Cell Analyzer (BD Biosciences, Heidelberg, Germany). Analysis was performed with FlowJo V10 software (BD Biosciences, Heidelberg, Germany). Cell surface expression is depicted as geometric mean of fluorescence intensity mediated by APC-coupled secondary antibody. Following primary antibodies with corresponding dilutions in FC buffer were used: α-HA (1:500, 901502, BioLegend, San Diego, US) and α-myc (1:1000, ab32, Abcam, Cambridge, UK). Secondary antibody used was α-mouse-APC diluted 1:200 in FC buffer (115-135-164, Jackson ImmunoResearch, West Grove, Pennsylvania, USA).

### 7. Statistical analysis

Statistical analysis was performed as described before (Düsterhöft et al 2021). In short, all quantified experiments were conducted at least three times and quantitative data is depicted as individual values with mean and standard deviation. Data was statistically analysed using general mixed model PROC GLIMMIX program (SAS 9.4, SAS-Institute, Cary, US) and tested for parametricity with residual analysis and Shapiro-Wilk test assuming normal, lognormal, exponential, binomial or logarithmic distribution. In case of heteroscedasticity (according to the covtest statement) degrees of freedom were corrected using Kenward-Roger approximation. Parametric data was analysed using one way or two way-ANOVA while for non-parametric data Man-Whitney test was applied using Graph Pad Prism 9.3.1 (GraphPad Software, San Diego, US). All p-values were corrected for multiple comparison using false discovery rate. Statistical significance was defined as p < 0.05 with */# p < 0.05 and **/## p < 0.01.

## Results

### The efficiency of soluble ACE2 release is influenced by natural occurring point mutations within the ACE2 stalk region

With the emergence of SARS-CoV-2 and the COVID-19 pandemic, several naturally occurring ACE2 point mutations have drawn increased attention, as they may influence viral entry and host response (Chen et al., 2021, Vadgama et al., 2022). Some of these variants appear to be enriched in specific populations and may contribute to observed differences in infection rates and disease severity across populations (Cao et al., 2020, Khayat et al., 2020, Vadgama et al., 2022). Three of the point mutations (N720D, N720S and L731F) are located within the ACE2 stalk region, an unstructured and flexible chain of 20 amino acids length connecting the extracellular collectrin-like and metalloprotease domain to the transmembrane helix (Fig. 1A-B). We have previously described that ACE2 shedding mainly occurs within the collectrin-like part by the sheddases ADAM10 and ADAM17 (Niehues et al., 2022). Furthermore, sheddase-mediated cleavage typically takes place in the juxtamembranous region.

**Figure 1:**
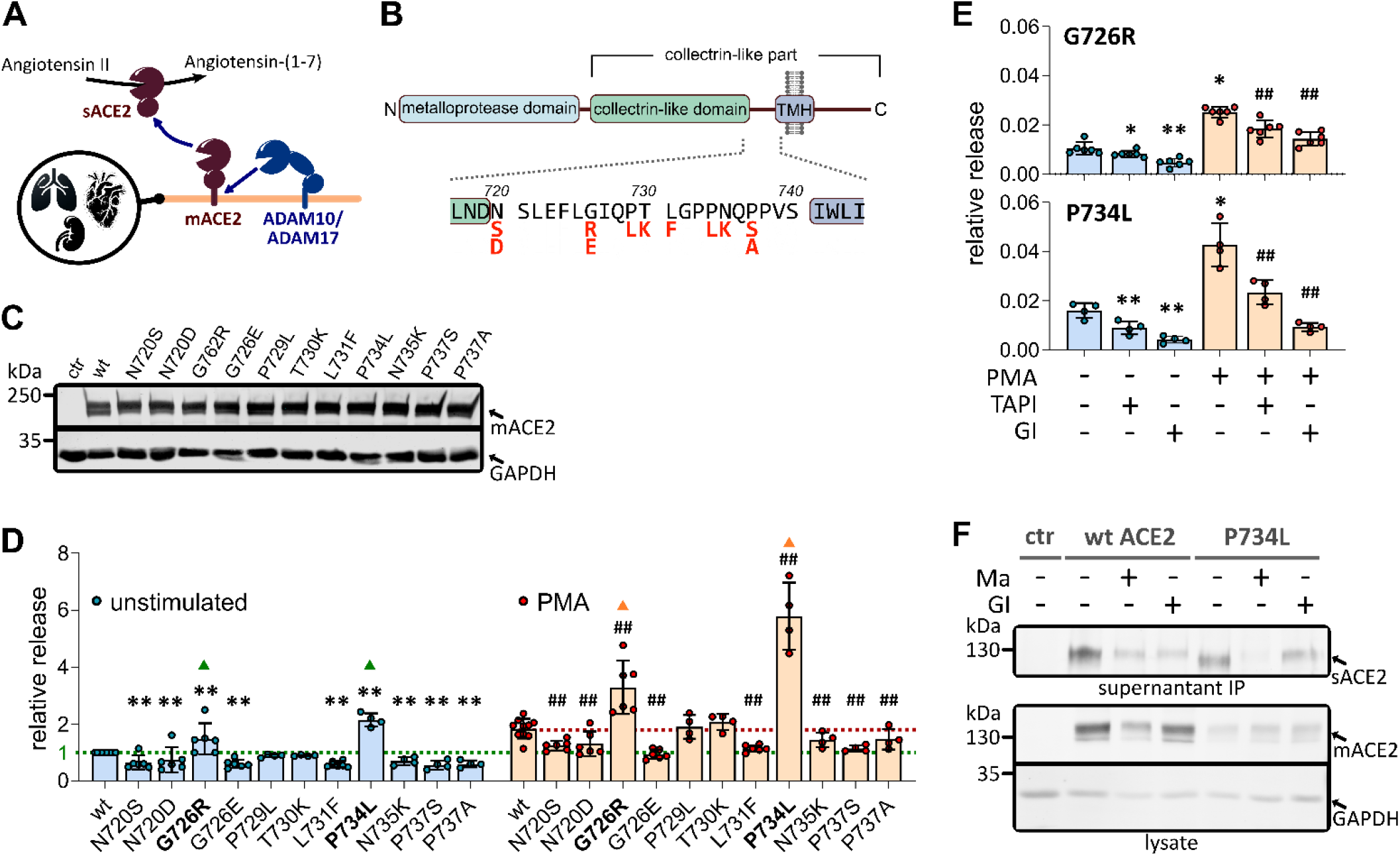
Natural occurring point mutations in ACE2 stalk region impact ACE2 release. **A)** ADAM10 and ADAM17 are the main sheddases responsible for the shedding of ACE2. ACE2 is expressed in the kidneys, lungs, and heart. **B)** ACE2 domain structure with detailed view of the stalk region (position 720 – 740). The naturally occurring point mutations in the stalk region are shown in red. N = N-terminus, C = C-terminus, TMH = transmembrane helix. **C)** HEK293 cells were transiently transfected with ACE2 variants tagged with alkaline phosphatase (AP). Western blot of inputs shows cell-associated/membrane-bound ACE2 variants (mACE2). **D)** Cells from C) were either left untreated or stimulated with 100 nm PMA for 24 hours. The relative release of ACE2 was assessed by calculating the ratio of the supernatant AP activity to the total AP activity (supernatant plus lysate). n > 3. **E)** HEK293 cells that transiently express the ACE2 variants ACE2_G726R or ACE2_P734L (tagged with AP) were either left untreated, or stimulated with 100 nM PMA, and/or treated with either 10 µM TAPI or 10 µM GI, for 24 hours. n > 3 **F)** HEK293 cells stably expressing myc-tagged ACE2 or the myc-tagged ACE2 variant ACE2_P734L were either left untreated or treated with 40 µM Marimastat (Ma) or 10 µM GI for 48 hours. Supernatants were used to enrich soluble myc-tagged ACE2 (sACE2) variants by coimmunoprecipitation. n = 2. **p-values)** to wt unstimulated: * < 0.05; ** < 0.01; to wt PMA: # < 0.05; ## < 0.01.

To assess how variation in the ACE2 stalk affects the release of soluble ACE2, we measured the release of eleven naturally occurring stalk point mutants (see Tab. 1 and Fig. 1B) using an N-terminal alkaline phosphatase (AP) tag. All variants were expressed in HEK293 cells as confirmed by western blotting (Fig. 1C). In the enzymatic shedding assay the release of soluble variants was measured in relation to the total cellular expression of each respective ACE2 mutant. Under unstimulated conditions, ACE2_G726R and ACE2_P734L exhibited significantly increased relative release, whereas most of the other mutants exhibited significantly less release than the wild type (Fig. 1D). ACE2_P729L and ACE2_T730K exhibited no change. Stimulation with the PKC activator PMA (phorbol 12-myristate 13-acetate), which is thought to predominantly promote ADAM17-dependent shedding, increased the release of all variants. ACE2_G726R and ACE2_P734L again showed significantly higher release, while most of the other mutants still showed reduced release compared to the wild type. As before, ACE2_P729L and ACE2_T730K show no difference (Fig. 1D).

To determine whether the increased release of ACE2_G726R and ACE2_P734L reflects enhanced susceptibility to either ADAM10 or ADAM17, we used TAPI (predominantly an ADAM17 inhibitor with additional activity against ADAM10 and other metalloproteases) and GI254023X (GI) (predominantly a potent ADAM10 inhibitor with 100-fold less potency for ADAM17). The release of both mutants was significantly inhibited by GI and TAPI, with ACE2_P734L showing a more pronounced sensitivity (Fig. 1E). The observation that GI was even more effective than GW in preventing constitutive and PMA-induced ACE2 release suggests that ADAM10 is the predominant sheddase in both conditions for both variants. This implies that the increased shedding of ACE2_P734L is to a large extent due to increased sensitivity to ADAM10 rather than ADAM17. Soluble wt ACE2 and ACE2_P734L were detected by co-immunoprecipitation in supernatants from cells stably expressing wt ACE2 or ACE2_P734L, respectively (Fig. 1F). In line with the results above, the soluble levels of both were reduced by GI and marimastat (Ma), which is a pan-metalloprotease inhibitor. Notably, in these stable cell lines, the abundance of cell-associated ACE2_P734L protein was markedly lower than that of wt ACE2.

Overall, most naturally occurring stalk variants reduced the release of ACE2, whereas the G726R and P734L mutations resulted in increased release. ACE2_P734L is particularly subject to ADAM10 -dependent shedding. However, release was not fully abolished by metalloprotease inhibition, consistent with the existence of additional metalloprotease-independent release routes, including extracellular vesicles, as we described earlier (Niehues et al., 2022).

### ACE2 mutant P734L shows elevated surface expression but reduced total ACE2 protein level

Stable expression of ACE2_P734L results in significantly increased release by shedding and reduced total cell-associated ACE2_P734L compared with wt ACE2, consistent with depletion through enhanced shedding. We therefore also quantified cell-surface ACE2 by flow cytometry. Unexpectedly, ACE2_P734L displayed significantly higher surface expression than wt ACE2 (Fig. 2A). Western blot analysis of matched lysates from the same samples again confirmed significantly lower total cellular ACE2_P734L (Fig. 2B,C). Consequently, ACE2_P734L showed an increased surface-to-total ratio (Fig. 2D), indicating preferential surface localisation and putative increased substrate availability. Thus, despite low total abundance, elevated surface expression likely contributes substantially to the enhanced shedding of ACE2_P734L.

**Figure 2:**
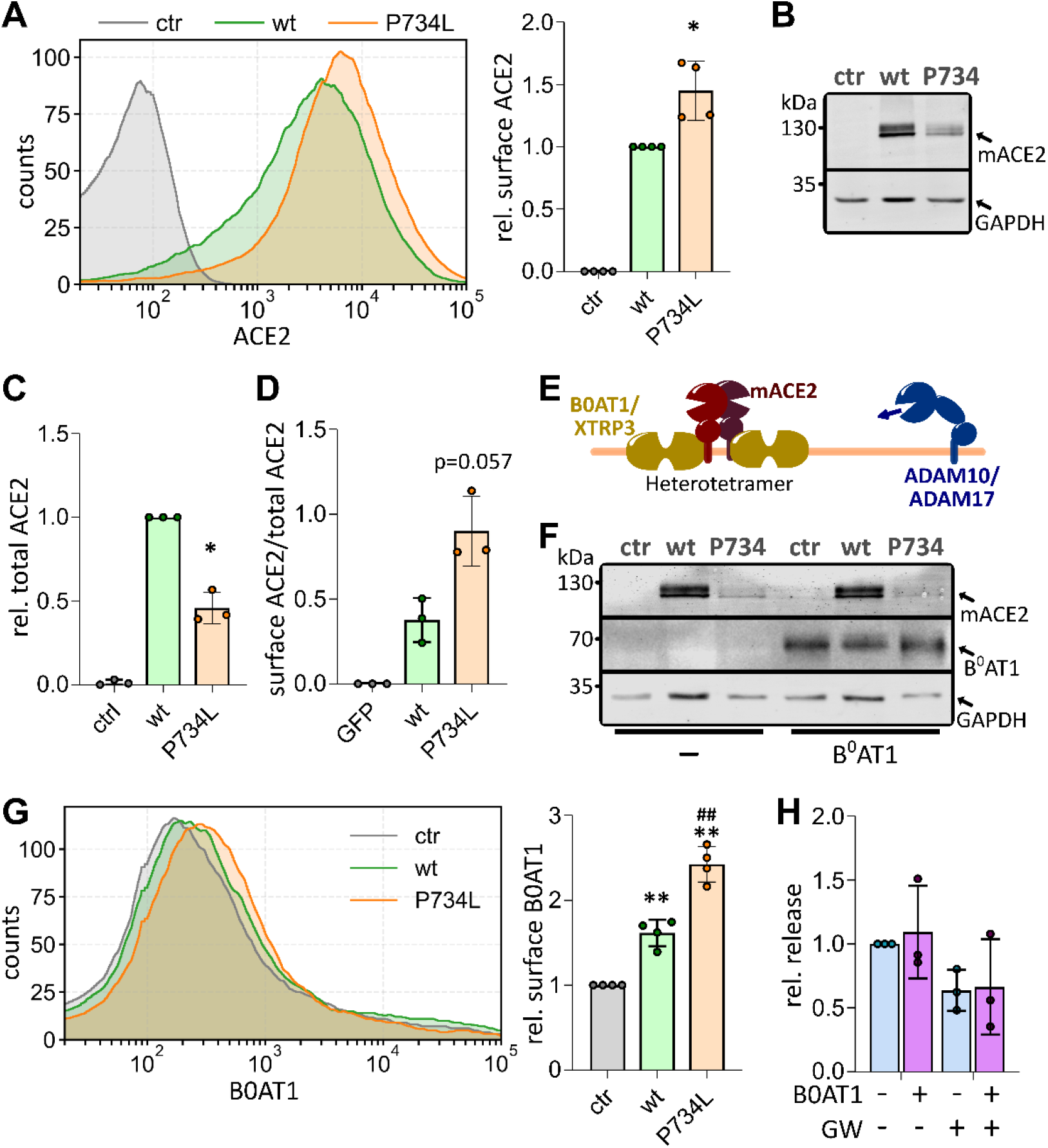
P734L mutation increases ACE2 and B0AT1 surface expression. **A-D)** HEK293 cells stably expressing wt ACE2, P734L, or GFP (control, ctr) were analysed by flow cytometry and Western blotting. Flow cytometry was used to quantify cell-surface abundance of the ACE2 variants (A), and Western blotting to assess total cellular abundance (B–C). The surface-to-total ACE2 ratio was then calculated (D). n > 3. **E)** Membrane-bound ACE2 (mACE2) forms homodimers and can assemble into heterotetrameric complexes with the amino acid transporters B0AT1 or XT3. This interaction is crucial for efficient cell-surface expression of the transporters. **F-G)** HEK293 cells stably overexpressing wt ACE2, P734L and or GFP (control, ctr) were transiently co-transfected with HA-tagged B0AT1 and subjected to Western blotting (F) and flow cytometric analysis (G). n = 4. **H)** AP-based shedding assay showing relative ACE2 release in the presence or absence of B0AT1 and upon inhibition with GW or GI for 24 h. n = 3. **p-values)** to wt: * < 0.05; ** < 0.01.

### P734L increases cell surface expression of the amino acid transporter B0AT1

The surface expression of the neutral amino acid transporter B0AT1 is known to depend on its interaction with ACE2 to form a heterotetramer (Fig. 2E). In line with earlier reports, indeed, we found that the additional expression of wt ACE2 significantly increases the surface abundance of B0AT1 (Fig. 2F,G). Next, we tested whether the higher ACE2_P734L surface localisation also increases B0AT1, even though ACE2_P734L is shed more extensively. Notably, we detected significantly higher levels of B0AT1 surface expression with ACE2_P734L compared to wt ACE2 (Fig. 2F,G).

As B0AT1 binds to the collectrin-like domain of ACE2 (Fig. 2E), which is targeted by sheddases, it was hypothesised that the formation of this complex could sterically restrict sheddase access, thereby modulating the release of ACE2. To test this, we performed a shedding assay in HEK293 cells that overexpress B0AT1. Interestingly, B0AT1 co-expression did not significantly alter ACE2 release compared to the control group (Fig. 2H), and the inhibitory effects of GW and GI were similar in both groups (with and without B0AT1). This suggests that shedding does not significantly impact the overall release of ACE2.

In summary, the ACE2_P734L mutation enhances B0AT1 surface expression and B0AT1–ACE2 complex formation does not measurably restrict protease access to ACE2 or alter ACE2 shedding under these conditions.

### Different ectodomain regions determine ACE2 shedding efficiency

ACE2 belongs to the ACE2 family, alongside ACE (ACE1) and TMEM27/collectrin, as it is an evolutionary chimera combining an ACE1-like metalloprotease domain with a TMEM27-like collectrin region (Fig. 3A). Notably, these related proteins exhibit distinct shedding behaviour: ACE1 is predominantly shed by ADAM10 rather than ADAM17, whereas TMEM27 is not efficiently shed by either enzyme (Fig. 3A).

**Figure 3:**
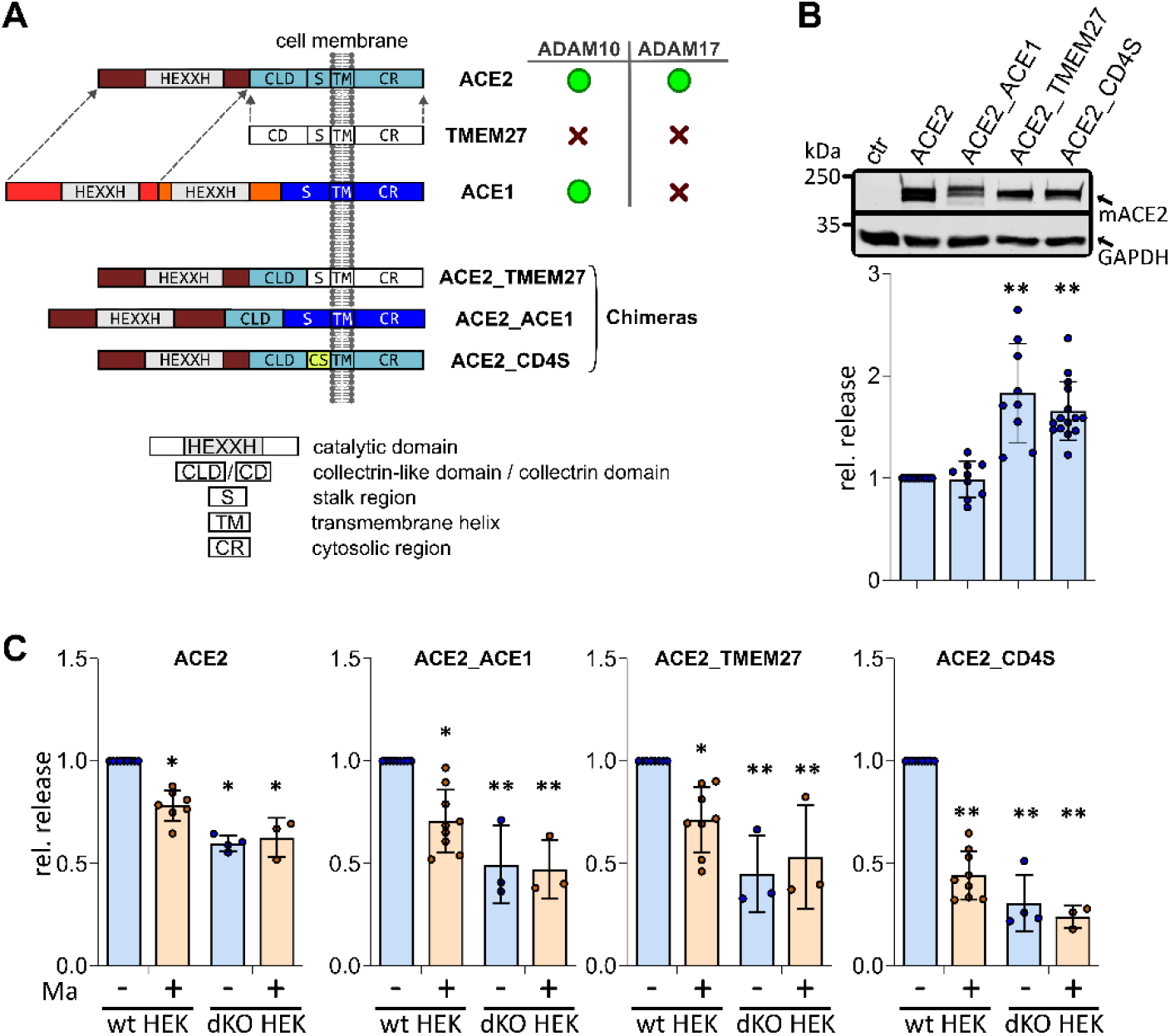
Different structural regions in members of the ACE2 family determiner shedding efficiency. **A)** Structural overview of ACE2 family members and their sensitivity to ADAM10 and ADAM17. ACE2 is an evolutional chimera from ACE and TMEM27 (collectrin). Furthermore, three ACE2 chimeras are shown. **B)** AP-tagged ACE2 chimeras and wt ACE2 were transiently overexpressed in HEK293 cells and incubated for 24 hours. Release of soluble proteins was quantified as before via AP-activity measuring. Western blotting shows equal expression. n > 9. **C)** AP-tagged ACE2 chimeras and wt ACE2 were transiently overexpressed in HEK293 or in ADAM10/ADAM17 deficient HEK293 cells (dKO HEK) and incubated for 24 hours either with or without marimastat (Ma). **p-values)** to wt: * < 0.05; ** < 0.01.

To investigate the role of the ACE2 stalk region for shedding efficiency, we created two chimeras (ACE2_ACE1 and ACE2_TMEM27), replacing the stalk, transmembrane segment and cytosolic tail with the corresponding regions of ACE1 or TMEM27. Shedding assays revealed that ACE2_ACE1 was released at levels comparable to wt ACE2 (Fig. 3B). Consistently, marimastat treatment and analysis in ADAM10/ADAM17 knockout cells revealed no difference from wt ACE2, indicating unchanged ADAM-dependent shedding sensitivity for this chimera (Fig. 3C).

As TMEM27 is not efficiently shed by ADAM10 or ADAM17, we expected ACE2_TMEM27 to be relatively resistant to shedding. However, we observed significantly increased release of ACE2_TMEM27 compared with wt ACE2 (Fig. 3B). Furthermore, treatment with marimastat and analysis in ADAM10/ADAM17 knockout cells significantly reduced its release, demonstrating that ACE2_TMEM27 acquired sensitivity to shedding by ADAMs (Fig. 3C).

A third chimera (ACE2_CD4S) had only its stalk replaced with that of CD4, an unrelated protein which has been reported not to undergo metalloprotease-related shedding. Nevertheless, like ACE2_TMEM27, ACE2_CD4S exhibited significantly increased release compared with wt ACE2, and this increase was predominantly ADAM-dependent shedding (Fig. 3B,C).

Overall, stalk point mutations and stalk replacements can markedly alter ACE2 shedding. However, their effects are context-dependent, indicating that determinants outside the stalk region also influence shedding efficiency.

## Discussion

ACE2 is a key regulatory component of the renin–angiotensin–aldosterone system (RAAS), balancing the vasoconstrictive and pro-inflammatory Ang II–AT1R axis on the one hand with the vasodilatory and anti-inflammatory Ang-(1–7)–MASR axis on the other. Similar to the related proteins ACE1 and TMEM27 (collectrin), ACE2 can be released as a soluble form (sACE2) through ectodomain shedding mediated by ADAM17 and ADAM10 (Lambert et al., 2005, Niehues et al., 2022). This study explores the role of the membrane-proximal ACE2 stalk region, which connects the extracellular domains to the transmembrane helix, in influencing shedding efficiency. Figure 5 provides an overview of the ACE2 constructs that were investigated and their respective constitutive release rates. The introduction of naturally occurring single point mutations (Tab. 1) and the use of ACE2 chimeras both had a significant impact on shedding rates. While no relevant change was observed for the ACE2_ACE1 chimera and the P729L and T730K mutants, most other ACE2 constructs exhibited a significant decrease in release rate compared to wt ACE2 (N720S, N720D, G726E, P729L, T730K, L731F, N735K, P737S, P737A), or an increase (G726R, ACE2_P734L, ACE2_TMEM27, ACE2_CD4S).

Importantly, neither ADAM10 nor ADAM17 recognizes a single, conserved cleavage motif in their substrates. Instead, substrate recognition and cleavage depends on a combination of different structural features of the ADAMs and their regulatory interactors such as iRhoms and tetraspanins (Zunke and Rose-John, 2017, Düsterhöft et al., 2019, Lipper et al., 2023), although certain amino acid patterns occur more frequently at cleavage sites than others (Tucher et al., 2014). These patterns reveal a common cleavage mechanism for ADAM10 and ADAM17 substrates, characterised by the presence of small and aliphatic residues before the cleavage site and a mostly hydrophobic environment on the prime side. The residue alanine (A) appears to be highly enriched at cleavage site position P2 and P1, whereas leucin (L) is the most prevalent residue at P1′. Furthermore, proline (P) appears to be enriched at P1 and other positions. Interestingly, arginine (R) also seems to be favoured at P1 or P2. In contrast, other charged and highly polar residues (notably D, E, K and N) are highly disfavoured around the cleavage site. This is consistent with the fact that ADAM cleavage favours flexible, low-charge segments.

Taking this into account, the wt ACE2 stalk/juxtamembrane region already exhibits a potential ADAM substrate environment. The membrane-proximal stalk contains a proline-rich segment (…LGPPNQPPVS…) immediately preceding the transmembrane helix, as well as a more hydrophobic patch (…FLGIQP…) further away (Fig. 4). These provide multiple potential cleavage targets. However, wt ACE2 does not appear to be maximally optimised for a single dominant motif overall, suggesting potential for improvement in the local P1/P1′ environment to enhance shedding efficiency. Applying the motif preferences to the naturally occurring single-point mutations suggests that altered shedding may reflect the optimisation or disruption of existing cleavage patterns or the creation of alternative cleavage sites. ACE2_P734L, the strongest “hypershedding” mutant, introduces a leucine within the proline cluster, a residue that is strongly favoured at P1′ (Tucher et al., 2014). This plausibly increases the efficiency of cleavage. Conversely, mutations that introduce charge (e.g. G726E and N735K) or remove proline from the proline-rich region (e.g. P737S and P737A) are consistent with the disruption of ADAM-compatible sequence environments and seem therefore aligned with reduced shedding. The different outcomes for mutations at the same position (G726R increases shedding, whereas G726E decreases it) further support the idea that local substitutions can either enhance or impair the “fitness” of cleavage sites, depending on whether they introduce residues that are compatible with the observed ADAM preferences (Tucher et al., 2014). The chimeras only partially support the conclusion that the stalk alone determines shedding efficiency. Importantly, the usage of ADAMs remains largely unchanged. Replacing the stalk/TM/cytosolic region with ACE1-derived sequences produced a release pattern similar to that of the wild type, whereas substituting it with TMEM27-derived sequences markedly increased release, which remained predominantly ADAM-dependent. Indeed, putative favoured cleavage sites for ADAM10 and ADAM17 exist in both the stalk region of ACE1 and the TMEM27 region (Fig. 4).

**Figure 4:**
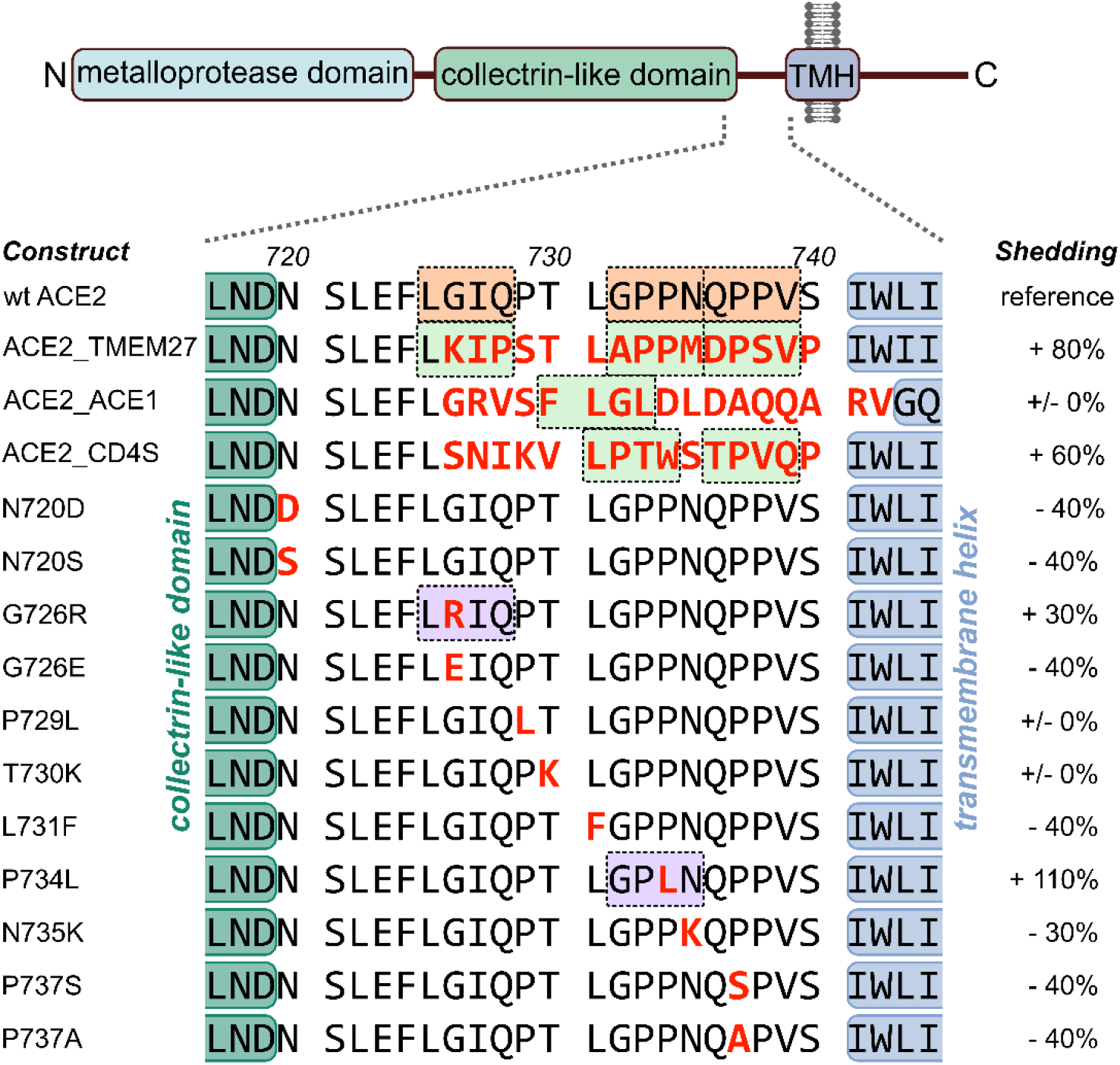
Stalk sequence and release rates of ACE2 mutants and ACE2 chimeras. Schematic overview of ACE2 with detailed view of the stalk region (position 720 – 740). Variant point mutations and ACE2 chimeras are highlighted in red. The difference in constitutive release compared to wt ACE2 is shown. The boxes indicate the potential ADAM10/ADAM17 cleavage sites.

Interestingly, under stable and lower (and therefore more physiological) expression levels, ACE2_P734L appears to be more likely to be transported to the cell surface. Combined with the increased shedding, this appears to result in the rapid depletion of total cellular ACE2_P734L compared to wt ACE2. However, the P734L single-point mutation in the ACE2 stalk also affects surface expression, which must be addressed in future studies.

A notable finding is the discrepancy between native TMEM27 and the ACE2_TMEM27 chimera. This shows that a simple “stalk alone determines shedding efficiency” model is not sufficient, but instead a multifactorial substrate recognition exists. While the stalk contributes to cleavage efficiency, several additional structural and regulatory layers are already known to influence ADAM substrate selectivity including (a) intracellular regulation via the cytosolic tail, (b) contributions from other ectodomain regions (e.g. local folding, glycosylation and domain–domain geometry), (c) the transmembrane helix as a potential recognition and (d) the function of ADAM regulator proteins that control substrate preference, most notably iRhom proteins for ADAM17 and TspanC8 tetraspanins for ADAM10 (Endres and Deller, 2017, Matthews et al., 2017, Düsterhöft et al., 2019). Furthermore, cleavage events outside the stalk region may contribute to the cleavage of ACE2. For example, TMPRSS2 cleaves ACE2 within a loop of the collectrin-like domain; however, this cleavage does not appear to generate soluble ACE2 (Heurich et al., 2014). Additionally, a putative ADAM17 cleavage site has been identified within the collectrin-like domain. However, this site is located within a more structured region and has not yet been fully validated (Heurich et al., 2014).

Overall, our data establish the stalk region as a major determinant of shedding efficiency, with naturally occurring variants markedly altering soluble ACE2 release. Further research is needed to identify the precise cleavage sites and to determine the relative contribution of stalk versus non-stalk shedding regulation.

Interestingly, the fusion protein ACE2_CD4S was also released via shedding. Earlier studies have described CD4 as not being subject to ectodomain shedding (Sadhukhan et al., 1998). However, our study revealed that not only was ACE2_CD4S released in the supernatant, but it was also released at significantly higher levels than wt ACE2. Interestingly, an earlier study proposed matrix metalloproteases as putative CD4 sheddases (Tseng et al., 2013). Nevertheless, our pharmacological and genetic inhibition of ADAM10/ADAM17 via marimastat or ADAM17/ADAM10 knockout respectively, significantly blocked the release of ACE2_CD4S. Furthermore, the CD4 stalk harbours putative ADAM10/ADAM17 cleavage sites (Fig. 4). Further studies are needed to clarify whether CD4 is shed by ADAMs, and the potential physiological and pharmacological impacts of this on the immune system and CD4-dependent HIV infection.

Our results indicate that single point mutations of ACE2 stalk region notably affect ACE2 release. While most mutants shed significantly less wt ACE2, the ACE2_G726R and ACE2_P734L mutants were released at levels 30% and 110% higher, respectively. This raises the question of whether individuals carrying these variants exhibit differences in SARS-CoV-2 susceptibility and COVID-19 disease course. Although the release/shedding of ACE2 is often referred to as a protective mechanism due to the decrease in the levels of the ACE2 receptor on the cell surface, and the resulting possible formation of soluble ACE2 scavenger receptors (Chan et al., 2020, Cocozza et al., 2020, Zoufaly et al., 2020), other studies have reported negative effects on the clinical course, as well as sACE2-mediated facilitation of SARS-CoV-2 infection (Rahman et al., 2021, Wang et al., 2021, Yeung et al., 2023). Some ACE2 variants are more frequent in distinct populations, giving a possible explanation for global differences in clinical courses (Cao et al., 2020, Khayat et al., 2020). Two of them, ACE2_N720D and ACE2_L731F, are part of the stalk region and were shown in this study to be released 40% less than wt ACE2. The ACE2_N720D variant is predominantly found in European populations, whereas the L731 variant is almost exclusively found in African populations. (Cao et al., 2020, Khayat et al., 2020). Altered shedding rates of the investigated variants could also be of great relevance for the RAAS, given that sACE2 levels were found to be elevated in patients with chronic kidney and cardiovascular diseases (Epelman et al., 2008, Anguiano et al., 2017). These elevated levels could be due to compensatory upregulation of the protective enzyme ACE2 or increased shedding rates mediated by inflammatory and hypoxic stimuli, as is known for other ADAM substrates such as ACE1 (Saftig and Reiss, 2011, Düsterhöft et al., 2019, Yu et al., 2026). Notably, there is a close association between the RAAS and SARS-CoV-2 infections, as the course of the disease seems to be characterised by RAAS imbalance (Elshafei et al., 2021, Rysz et al., 2021). Further clinical and genetic studies are needed to investigate possible correlations between these factors and clinical parameters of SARS-CoV-2 infection and cardiovascular diseases.

Another important function of ACE2, in which its shedding may play a role, is binding to amino acid transporters such as B0AT1. ACE2 is essential for the expression of B0AT1 on the surface of cells, and thus for the uptake of amino acids in the small intestine and other tissues (Camargo et al., 2020). In our study, we also found that B0AT1 expression levels were higher when ACE2 expression levels were increased. Notably, we showed that the ACE2_P734L variant itself has higher surface expression and can thereby also promote higher B0AT1 surface expression compared to wt ACE2. Previous studies have reported that B0AT1/ACE2 heterodimers form tetramers, and that this interaction is mediated by the collectrin-derived parts of ACE2 (Yan et al., 2020, Stevens et al., 2021). This could potentially hinder sheddase access to the ACE2 cleavage site, as indicated by molecular docking modelling (Andring et al., 2020, Stevens, 2020). In contrast, our shedding assays revealed that the presence and absence of B0AT1 does not alter ACE2 release, indicating that complex formation is subordinate to sheddase recognition and the shedding process.

Overall, our data highlights the importance of the ACE2 stalk region in shedding. Several natural variants have been shown to be released to varying degrees, which may affect RAAS homeostasis, SARS-CoV-2 infection and functions of amino acid transporters such as B0AT1.

## Abbreviations

ACE2: angiotensin-converting enzyme 2
AP: alkaline phosphatase
BSA: bovine serum albumin
DMSO: dimethyl sulfoxide
FCS: fetal calve serum
GI: GI254023X
GW: GW280264X
HEK: human embryonic kidney cells
mACE2: membrane-bound ACE2
PBS: phosphate buffered saline
PCR: polymerase chain reaction
PNP: p-nitrophenol
PNPP: p-nitrophenyl phosphate
sACE2: soluble ACE2 (ectodomain)
TAPI-1: Tumor necrosis factor α protease inhibitor TBS tris buffered saline
wt: wildtype

